# Genetic basis of maneb-induced dopaminergic neurodegeneration in *Drosophila*

**DOI:** 10.1101/2025.05.07.652747

**Authors:** Stefanny Villalobos-Cantor, Alicia Arreola-Bustos, Ian Martin

## Abstract

Parkinson’s disease (PD) is a complex neurodegenerative disease driven by combined genetic and environmental factors. Human studies support increased PD risk following exposure to the pesticide maneb yet animal studies generally report subtle or no dopaminergic phenotypes unless maneb is combined with additional pesticides. Consequently, it is unclear whether exposure to maneb alone promotes dopamine (DA) neurodegeneration and if so, what the underlying mechanisms are. We hypothesized that gene-environment interactions are major determinants of maneb-mediated neurodegeneration and in support of this find that DA neuron viability is quantitatively divergent among 186 maneb-exposed genetically varying fly strains from the *Drosophila* Genetic Reference Panel (DGRP). Through genome-wide association analysis we identify several candidate genetic modifiers of maneb-induced DA neurodegeneration and further validate two candidate genes, *fz2* and *CG14186* which we find potentiate maneb-induced DA neurodegeneration when knocked-down. *fz2* and the mammalian ortholog of *CG14186* (*TMEM237*) are both thought to be necessary for intact Wnt pathway signaling in nervous system development and maintenance. Accordingly, we find that adult-specific perturbation of Wnt signaling is sufficient to promote maneb-induced DA neuron loss. Collectively, these results support a role for gene-environment interactions in PD etiology and reveal candidate mediators of maneb-related DA neurodegeneration *in vivo*.

**ARTICLE SUMMARY:** Exposure to the pesticide maneb has been linked to increased PD risk although animal models of neurodegeneration have produced mixed results. We developed a *Drosophil*a model for delayed onset dopamine neuron loss following maneb exposure and find that maneb-related neurodegeneration is quantitatively divergent across a large collection of *Drosophila* strains. We use a genome-wide association approach with follow-up validation to identify candidate modifier genes, uncovering roles for *fz2* and *CG14186*. Both genes are implicated in Wnt signaling and we further show that deregulated Wnt signaling in adults promotes maneb-induced dopamine neuron loss. These studies support a role for gene-environment interactions in maneb neurotoxicity and yield insight into the underlying genes involved.

## INTRODUCTION

Parkinson’s disease (PD) is a common movement disorder caused primarily by a loss of dopaminergic (DA) neurons in the substantia nigra. Epidemiological studies link pesticide exposure to elevated PD risk, yet human studies alone cannot conclusively link PD etiology to any single pesticide due to difficulties with exposure assessment (1). Controlled animal studies are therefore crucial for determining whether pesticide exposure results in PD-related neurodegeneration and if so, can be harnessed to probe mechanisms underlying disease development (2). For many pesticides including the fungicide maneb, a lack of robust neurodegenerative phenotypes has hindered advances in our biological understanding of disease development. One limitation of prior studies is a failure to consider gene-environment interactions that may be critical determinants of neurodegeneration. Thus, a lack of pesticide-induced neurodegeneration in a single isogenic animal strain may be specific to that strain and mask the neurodegenerative potential of a pesticide among a genetically varying population. Animal PD models that account for gene-environment interactions therefore offer a large untapped potential to provide insight into PD etiology which may be advanced to identify potential therapeutic targets.

Maneb (manganese ethylene-bis-dithiocarbamate) belongs to the dithiocarbamate class of fungicides that also includes zineb (zinc ethylene-bis-dithiocarbamate) (3). Maneb and mancozeb (maneb and zineb combined), are used throughout the U.S. and other countries on a wide range of fruits, vegetables, and field crops because they control a wide spectrum of fungal diseases (1, 2). Maneb is used to prevent crop damage in the field and to protect harvested crops from deterioration in storage or transport. Case reports of Parkinsonism in agricultural workers exposed to maneb (4–6) have generated interest in maneb as a possible PD-causing agent. Data from the Agricultural Health Study found an association of maneb/mancozeb with PD, although statistical power was limited by the small number of cases (7). Maneb is thought to cross the blood-brain barrier and while evidence from animal studies is mixed, there are reports suggesting that it is capable of exerting neurotoxic effects that may affect the dopaminergic system (3, 8–10). For example, intraperitoneal injection of maneb (30 mg/kg) in C57BL/6 mice twice per week for 6 weeks has been shown to cause locomotor activity deficits and affect levels of striatal DAT, dopamine and its metabolite DOPAC, but with no overt dopamine neuron loss (9, 10). Knowledge of the mechanism for maneb/mancozeb is limited as studies have generally targeted the analysis of mitochondrial or ubiquitin-proteasome system (UPS) function for potential perturbations without taking an unbiased approach. These studies either suggest (8, 11) or refute (12, 13) the idea that maneb may promote ROS production/oxidative stress and one study reported the potential for maneb injected intraventricularly in rats to produce complex III deficits in brain mitochondria tested *in vitro* (8). Maneb was reported to inhibit the UPS in SK-N-MC neuroblastoma cells or in a dopaminergic cell line (14). Aldehyde dehydrogenase (ALDH) activity, important in detoxifying the dopamine metabolite DOPAL has been linked with DA neuron toxicity, and maneb along with other dithiocarbamates reportedly inhibit ALDH activity *in vitro* (15). Interestingly, genetic variation in ALDH2 exacerbated PD risk in subjects exposed to pesticides that could inhibit ALDH suggesting that ALDH activity may be a target of maneb action, although this has not been verified *in vivo* (15).

Hence, subtle animal phenotypes and a generally restricted investigation into potential mechanisms of action has limited biological insight into maneb-related neurodegeneration. Consequently, the neurodegenerative potential for maneb and underlying mechanisms await clarification and could impact our understanding of PD development. Here, we used an unbiased genome-wide approach to probe the genetic basis of maneb-related DA neurodegeneration. We demonstrate that PD-related neurodegeneration is quantitatively divergent in exposed strains from the *Drosophila* Genetic Reference Panel (DGRP). This set of ∼200 fly strains have been inbred to homozygosity and fully sequenced, allowing the identification of loci associated with any trait of interest that can be assessed in each of the strains (16). We examined genome-wide association of single nucleotide polymorphisms (SNPs) with DA neuron viability across these strains and through follow-up functional testing uncovered *fz2* and *CG14186* as novel candidate modifier genes for maneb neurotoxicity. There is evidence to support a potential functional overlap between *fz2* and the mammalian ortholog of *CG14186* (*TMEM237*) in the conserved Wnt signaling pathway thus suggesting a role for deregulated Wnt signaling in maneb-related DA neurodegeneration.

## RESULTS

### A *Drosophila* model for maneb-induced neurodegeneration

Evidence from case reports and epidemiological studies indicate that PD associated with maneb or mancozeb exposure manifests years after initial exposure suggesting a potential lag between exposure and neurodegeneration (5, 7). Toward developing a suitable maneb exposure model, we sought a maneb concentration which when added to *Drosophila* food would (i) not modify food consumption rates, (ii) cause a delayed onset DA neurodegenerative phenotype and (iii) not cause severe lethality to avoid selective bias in flies assessed for neurodegenerative phenotypes. Using a concentration range informed by prior studies (17, 18) we observed that 1 mM maneb has no significant effect on fly feeding behavior while 10 mM significantly impairs feeding (Fig. 1A). We therefore decided to use 1 mM maneb in subsequent studies to avoid potential confounding effects of reduced feeding on neurodegeneration (19). When applied to two prototypic DGRP strains (RAL-911 and RAL-177), housing flies on 1 mM maneb-supplemented food for 7days results in no immediate DA neuron loss at the 7-day timepoint, while subsequently housing maneb exposed flies for an additional 14 days on standard food after maneb exposure leads to significant DA neuron loss in both strains (Fig. 1B), consistent with delayed onset neurodegeneration. The majority (65%) of maneb exposed flies remain alive at the end of this 21-day period for both DGRP strains, hence flies assessed for DA neuron viability are considered representative of their populations. Using this maneb concentration and timeline we were able to satisfy our model criteria, hence we used this approach on the wider DGRP strain collection.

**Figure 1.**
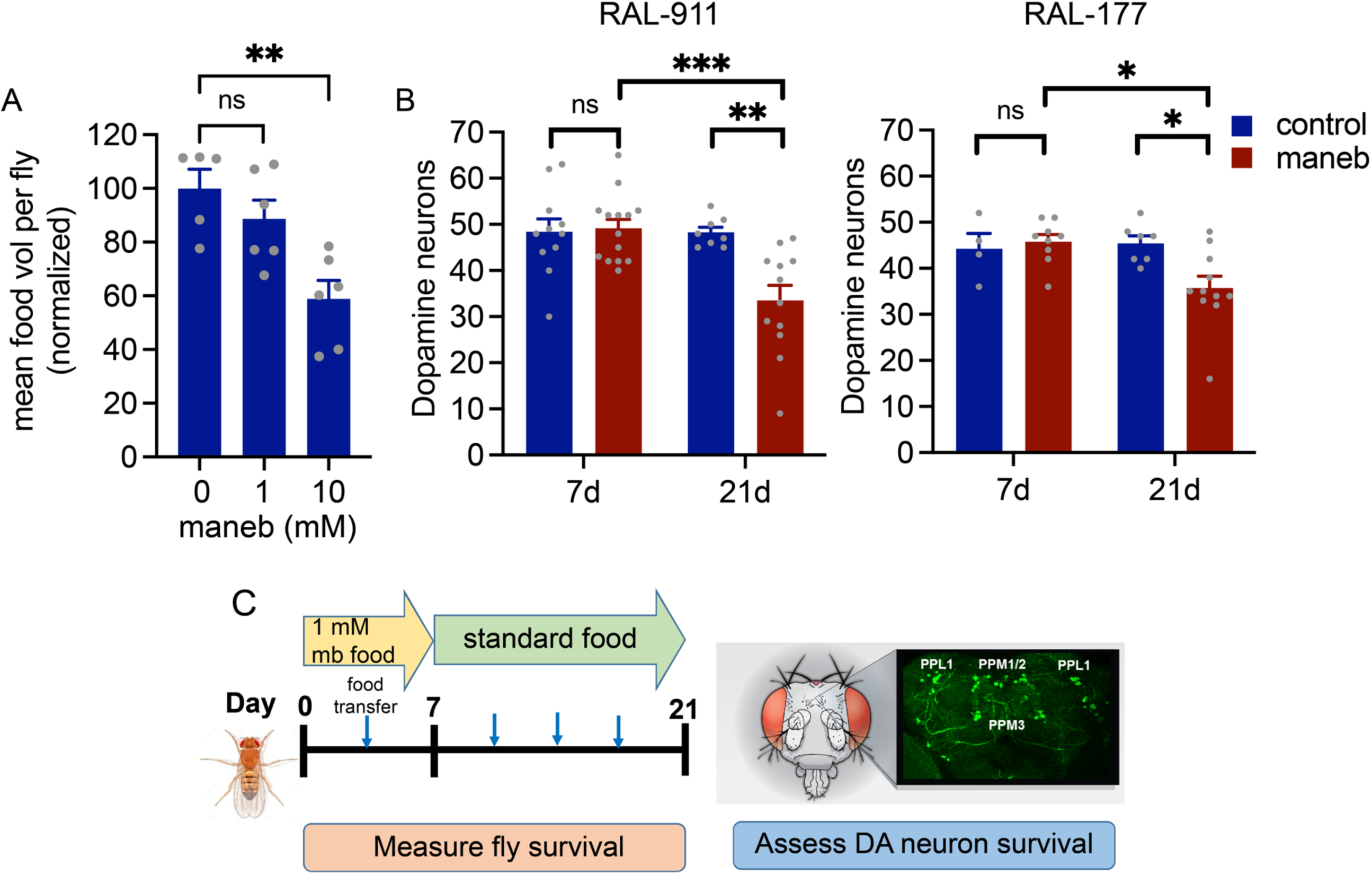
A model for delayed onset maneb-induced DA neurodegeneration in *Drosophila*. (A) Significant effect of 10 mM but not 1 mM maneb on feeding behavior of *w^1118^* flies (ANOVA, P <0.01, Dunnett’s post-hoc tests, ** P <0,01, n =5-6 groups of 8-10 flies per genotype). (B) flies belonging to RAL-911 and RAL-177 DGRP strains exhibit intact DA neuron counts (PPM1/2, PPM3 and PPL1 clusters combined) at 7 days of maneb exposure but significant DA neuron loss following an additional 14 days of aging on standard food (individual two-way ANOVAs for maneb and age, Šídák’s post-test * P <0.05, ** P <0.01, *** P <0.001, n =4-12 female brains per group). (C) scheme illustrating exposure to maneb-supplemented food and testing timeline.

### Maneb-induced DA neuron loss varies across the DGRP

We assessed DA neuron viability in 186 DGRP strains across 15 experimental batches (Methods) and observe substantial variation in batch-corrected DA neuron counts (ANOVA, *P* = 1.73×10^−170^) (Fig. 2A and Table S1). Using individual brain DA neuron counts from all maneb-exposed strains, we estimate a broad sense heritability for DA neuron viability of *H^2^*= 84.9% indicating that a large portion of phenotypic variation observed is attributable to genetic factors. Some strains retain maximal numbers of DA neurons at the end of the 21-day testing period while in other strains, DA neuron viability is substantially compromised (Fig. 2B). In lieu of assessing control (no maneb exposure) DA neuron integrity in all 186 DGRP strains, we attempted to probe the maneb-dependence of the DA neuron counts we obtained through a two-pronged approach. First, to determine whether low DA neuron counts are representative of maneb-induced neuron loss, we chose 20 DGRP strains exhibiting the lowest DA neuron counts of all 186 strains assessed, aged these flies on standard food medium without maneb for 21 days and assessed DA neurons as performed for maneb exposed flies. Comparison of maneb-exposed and unexposed groups for each of these 20 DGRP lines reveals that maneb causes reduced mean DA neuron numbers for all lines and that this reaches statistical significance in all but 5 lines (Fig. 2C), consistent with pervasive DA neuron loss. Second, we compared our measures of DA neuron viability across all 186 DGRP strains (Table S1) with published DA neuron counts obtained from the same DGRP strains simply aged for 21 days without any pesticide exposure (20).

**Figure 2.**
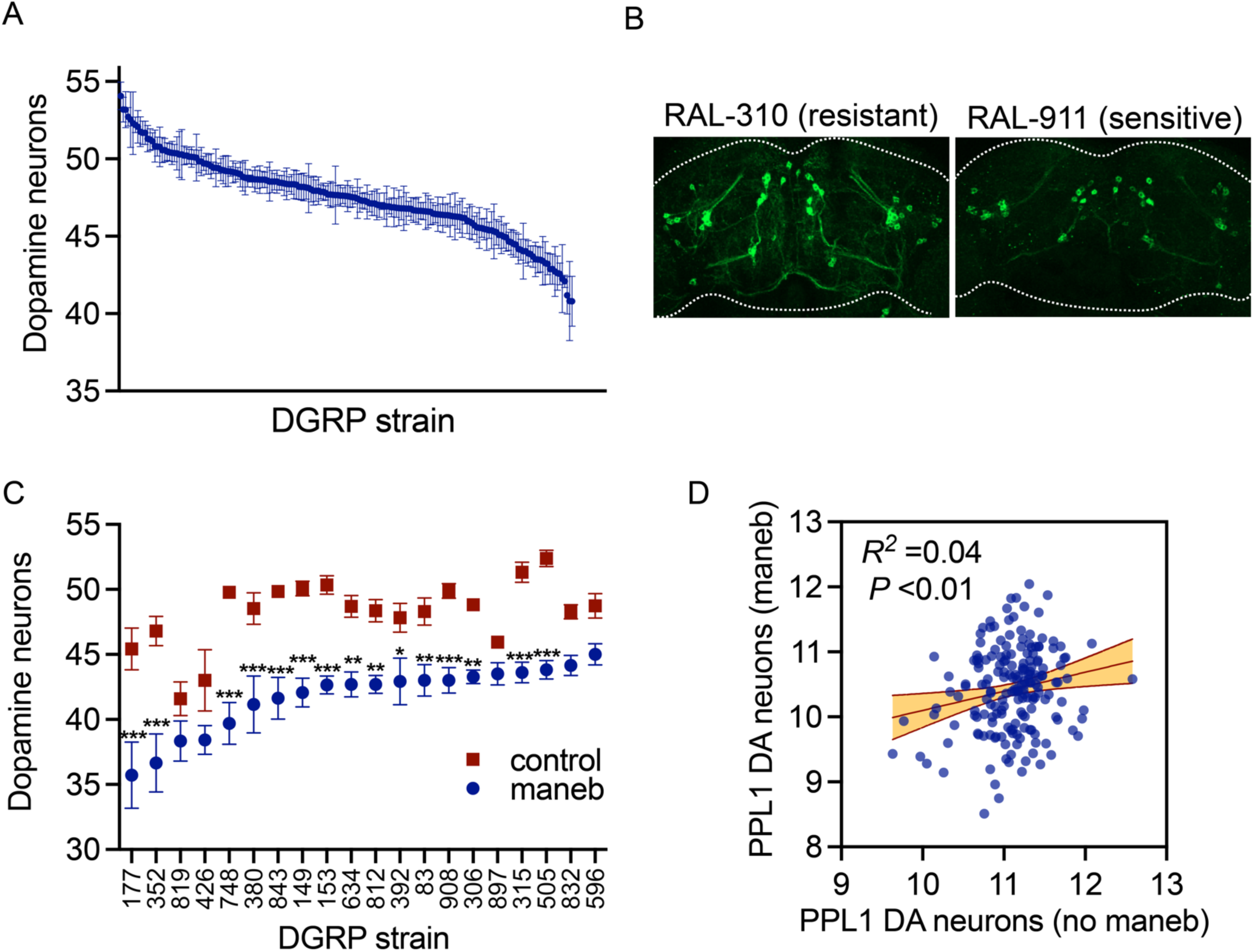
DA neuron viability divergence following maneb exposure. (A) Batch-corrected DA neuron counts for 186 DGRP strains (see Methods and Table S1 for individual DGRP line phenotypes) following 7 days of maneb exposure followed by 14 days aging. DGRP background has a strong effect on DA neuron counts (AVNOA *P* = 1.73 x 10^−170^, *n* =12-20 brains/genotype). (B) projection images of TH-immunostained brains from a maneb-resistant (RAL-310) and a maneb-sensitive (RAL-229) DGRP strain. (C) Variable DA neuron viability (not batch-corrected) at day 21 assessed via TH immunostaining in females of 20 DGRP lines fed vehicle-(control) or 1 mM maneb-supplemented food. Significant differences between the groups are indicated (2-way ANOVA for effect of strain (*P* < 0.0001) and treatment (*P* < 0.0001), interaction (*P* < 0.0001), Šídák’s post-test, * *p* < 0.05, ** *p* < 0.01, *** *p* < 0.001, *n* = 12-20 brains/group). (D) weak linear regression between PPL1 DA neuron counts in maneb-exposed flies and age-matched controls over measures from 186 DGRP strains. Age-matched control data are derived from Davis *et al*. (Ref 20). Best-fit line and shaded region is between 95% confidence bands.

This uncovered a very weak but significant regression coefficient (*R^2^* =0.04, *P* <0.01) for PPL1 DA neuron viability between the two datasets (Fig. 2D). Taken together, these data suggest that there may be a minor influence of maneb-independent factors such as strain-specific development or aging on our DA neuron counts from maneb-treated flies (Fig. 2A), but that maneb exposure appears to be the predominant driver of DA neuron viability divergence. Mean survival of all 186 lines housed on maneb food at the end of the 21-day testing period is 96% ± 0.6% (Table S1), hence, mortality levels are generally low at this restrictive maneb concentration and should not introduce major selective bias in the surviving flies examined for DA neurodegeneration.

### Genome-wide association to identify candidate maneb modifier genes

We assessed DA neuron counts across 186 DGRP strains (Table S1) for association with ∼1.8 million single nucleotide polymorphisms (SNPs) across the fly genome (Methods and Data S1). Analysis over this number of strains is underpowered to identify polymorphisms with genome-wide significance which was not the goal of this study. Instead, we used these SNP associations to nominate candidate modifier genes influencing maneb-induced DA neuron degeneration to advance for subsequent validation. We identified 146 SNPs in a total of 93 genes that associate with maneb-induced DA neuron death at a nominal *P*-value of *P* <10^−5^ (Table S2) of which 19 SNPs in 12 candidate genes are associated at a higher stringency threshold of *P* <10^−6^ (Table 1) and are considered top association candidates. These top association SNPs are distributed across the fly genome (Fig. 3A) and pairwise linkage disequilibrium (LD) analysis shows that they represent 14 independent loci (*r^2^* <0.5). SNP pairs in LD are all within the boundaries of the same gene suggesting that the overall impact of LD on our genome-wide association results is low (Fig. 3B). The majority of genes harboring these top association SNPs have putative human orthologs (Table 1) and represent a range of predicted biological functions including actin cytoskeleton regulation (*cv-c*, *cortactin* and *cdi*), genes annotated for a role in axon guidance (*Sema-2a*), ubiquitin ligase activity (*CG10283*), regulation of kinase signaling (*Ptp61F*), transmembrane transport (*CG7342*) and wnt signaling and cilium assembly (*CG14186*). Furthermore, gene ontology (GO) analysis for biological process enrichment in all SNPs associating at *P* <10^−5^ reveals an enrichment for cadherin signaling (14.7-fold enrichment, *P* =3.4×10^−5^, FDR =5.4×10^−5^) and Wnt signaling (7.1-fold enrichment, *P* =2.6×10^−4^, FDR =2.0×10^−4^) (Table 2). Unsurprisingly given the interconnectedness of Wnt and cadherin signaling pathways there is a major overlap in the genes annotated within these two enriched pathways (Table 2).

**Figure 3.**
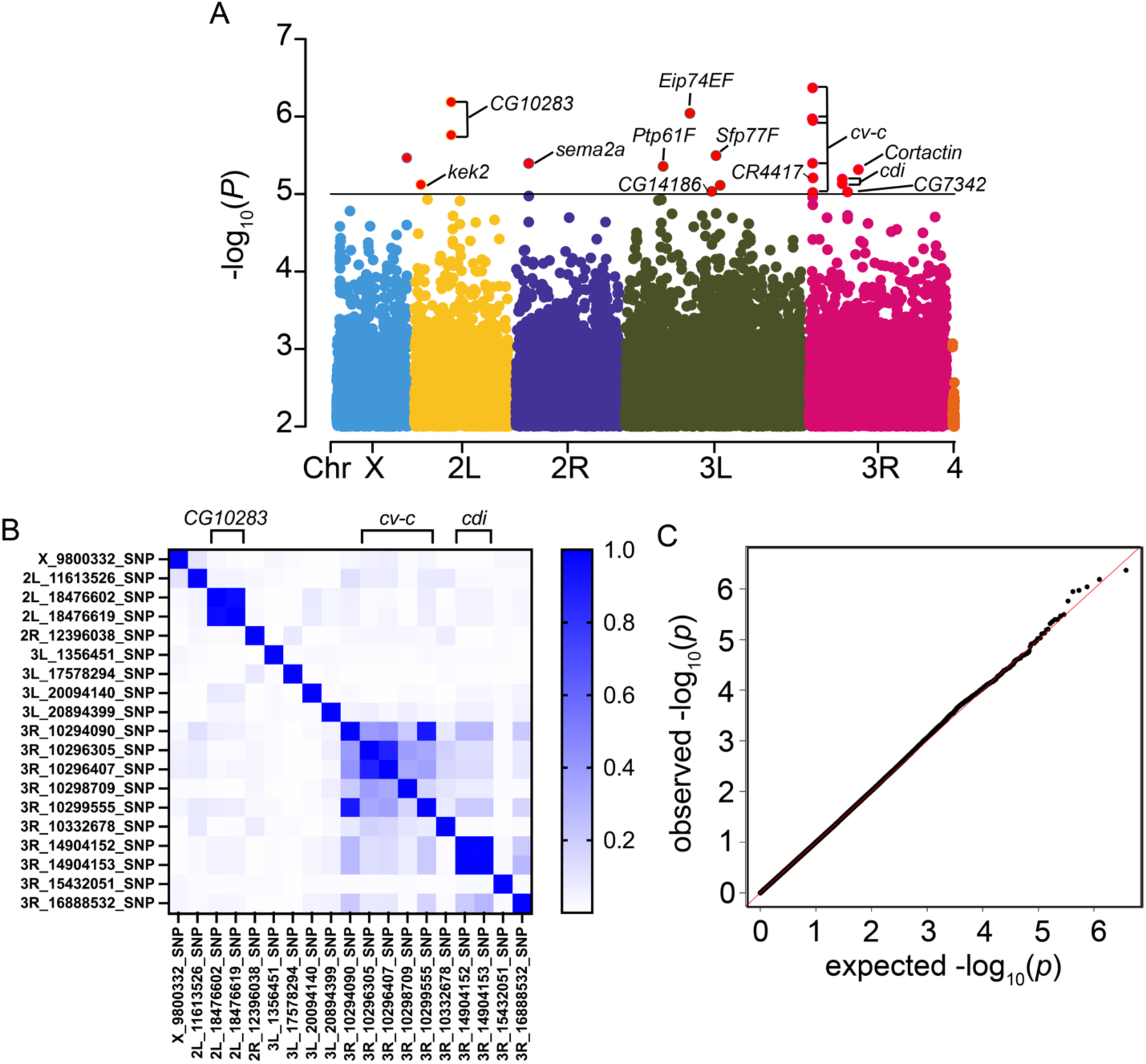
Genome-wide association for maneb-induced DA neurodegeneration. (A) Top association SNPs (Table 1 and Data S1) are distributed across the fly genome. – log_10_ *P*-values for each SNP (MAF ≥ 0.05, < 30% missingness) associating with DA neuron viability following maneb exposure (*P*-value cutoff of *P* <0.01). SNPs meeting the *P* <10^−6^ threshold are annotated with the genes harboring these SNPs unless they are intergenic in nature. (B) A pairwise LD heatmap for 19 top associating SNPs and 1 associating deletion (*P* <10^−6^), groups of SNPs in LD are indicated with brackets at the top and fall within genes *CG10283*, *cv-c* and *cdi*. A color scale for *r^2^* is shown. (C) A Q-Q plot of –log_10_ *P*-values for the variants included in the GWAS shows the data fit a normal distribution.

**Table 1.** Top association candidate genes Genes identified for all SNP associating with a linear mixed model-derived *P* <10^−6^. ^a^SNP with most statistically significant association for the candidate gene.

| Rank | SNP<sup>a</sup> | Candidate gene | P-value | Human ortholog |
| --- | --- | --- | --- | --- |
| 1 | 3R:10296407 | *cv-c* | 4.85E<sup>-07</sup> | *DLC1* |
| 2 | 2L:18476602 | *CG10283* | 6.49E<sup>-07</sup> | *CCDC50* |
| 3 | 3L:17578294 | *Eip74EF* | 9.08E<sup>-07</sup> | *ELF2* |
| 4 | 3L:20894399 | *Sfp77F* | 3.19E<sup>-06</sup> | — |
| 5 | 2R:12396038 | *Sema-2a* | 4.03E<sup>-06</sup> | *SEMA3A* |
| 6 | 3L:1356451 | *Ptp61F* | 4.39E<sup>-06</sup> | *PTPN1/2* |
| 7 | 3R:16888532 | *Cortactin* | 4.84E<sup>-06</sup> | *CTTN* |
| 8 | 3R:10332678 | *CR44177* | 6.19E<sup>-06</sup> | — |
| 9 | 3R:14904153 | *cdi* | 6.41E<sup>-06</sup> | *TESK2* |
| 10 | 2L:11613526 | *kek2* | 7.56E<sup>-06</sup> | *LRRC24* |
| 11 | 3L:20094140 | *CG14186* | 9.24E<sup>-06</sup> | *TMEM237* |
| 12 | 3R:15432051 | *CG7342* | 9.40E<sup>-06</sup> | *SLC22A23* |

**Table 2.** PANTHER pathway analysis. Enrichment analysis for 93 genes identified for SNP associating with a linear mixed model-derived *P* <10^−5^. Reference genome is the number of genes ascribed to a biological process in the entire *Drosophila* genome (*n* =13,767 genes). Genes is the number and names of associating genes in a pathway. Expected is the gene number expected in a pathway under the null hypothesis.

| PANTHER pathway | Ref. genome (D. mel) | Genes | Expected | Fold-enrichment | P-value | FDR |
| --- | --- | --- | --- | --- | --- | --- |
| Cadherin signaling | 48 | 5 (*Cad99C*, *CadN2*, *Cad87A*, *fz*, *fz2*) | 0.34 | 14.69 | 3.43E<sup>-05</sup> | 5.38<sup>-05</sup> |
| Wnt signaling | 120 | 6 (*Cad99C*, *CadN2*, *Cad87A*, *fz*, *fz2*, *Antp*) | 0.85 | 7.05 | 2.6E<sup>-04</sup> | 2.04<sup>-02</sup> |

### Functional analysis of nominated genes and pathways

We next carried out gene-level RNAi studies to identify candidate genes involved in resistance to maneb-mediated DA neurodegeneration, based on the premise that if a candidate gene is important in mediating maneb resistance then its silencing would promote maneb-induced DA neuron loss. We targeted all genes nominated at a *P* <10^−6^ SNP association threshold (Table 1) and all genes belonging to *cadherin signaling* and/or *Wnt signaling* uncovered by our GO analysis. RNAi lines were available for all but two of our top associating genes (*Cortactin* and *CR44177*), hence in total we performed RNAi on 16 individual genes (10 top association genes and 6 cadherin/Wnt signaling genes). We mated RNAi lines (or vector control line) for each candidate gene to flies harboring the *TH-GAL4* driver in a single experimental batch to drive DA neuron-specific gene silencing and assess cell autonomous effects on DA neuron viability. As with DGRP testing, we exposed flies to 1 mM maneb continuously for 7 days then housed them on standard food without maneb for an additional 14 days before assessing DA neurons. A genetic control strain harboring the RNAi vector does not exhibit significant DA neuron loss when exposed to maneb, while flies with *fz2* or *CG14186* knock-down in the same genetic background manifest significant maneb-induced DA neuron loss (Fig. 4A) suggesting a potentiation of maneb effects after silencing either gene. An enhancer effect of knocking-down these genes is consistent with their ability to contribute towards maneb resistance when their expression is intact. Additionally, RNAi to *Eip74EF, Sema-2a*, and *fz* each result in fewer DA neurons in the absence of maneb, suggesting that these genes may be required for the baseline development and/or maintenance of DA neuron integrity independent of maneb exposure (Fig. 4A). The size effect of individual gene knock-down on maneb-induced neurodegeneration appears low, which when taken together with the lack of overt DA neuron loss in most RNAi lines exposed to maneb supports the interpretation that maneb-mediated DA neuron loss may be highly polygenic in nature. For *fz2* and *CG14186* RNAi, we queried whether the DA neuron loss observed following maneb exposure was attributable to one or more individual DA neuron clusters. Analysis of DA neuron clusters indicates that *fz2* knock-down is associated with significant loss of PPL1 DA neurons, while DA neuron reductions within other clusters did not meet the threshold for significance. Conversely, CG*14186* knock-down results in significant loss of DA neurons in PPM1/2, PPM3 and PPL1 clusters upon maneb exposure (Fig. 4B), hence these data suggest that among the DA neuron clusters included in our analysis, maneb does not appear to exert its neurotoxic effects on any single cluster exclusively.

**Figure 4.**
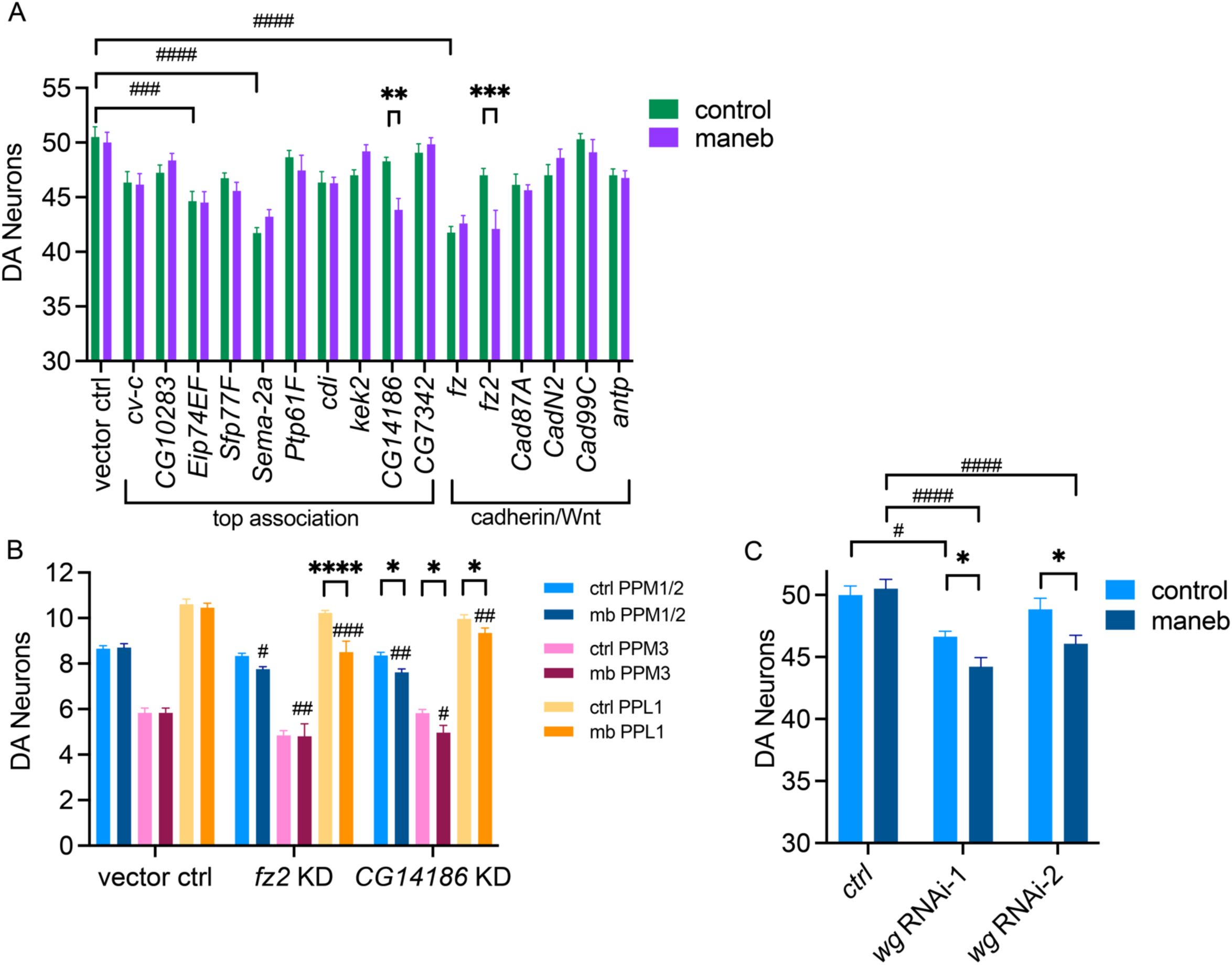
Gene-level testing implicates the Wnt signaling pathway. (A) DA neuron-specific knock-down (via *TH-GAL4*) in RNAi lines targeting 16 candidate genes or a vector control (Methods). *CG14186* or *fz2* knock-down results in significant DA neuron loss in maneb exposed flies (two-way ANOVA, Šídák’s post-test, ** P <0.01, *** P <0.001, n =12-20 brains per genotype) while DA-specific knock-down of *Eip74EF*, *Sema-2a* and *fz* impacts baseline DA neuron viability in controls (Šídák’s post-test, ### P <0.001, #### P <0.0001). (B) individual DA neuron cluster analysis reveals significant loss of PPL1 DA neurons for *fz2* and significant loss of PPM1/2, PPM3 and PPL1 DA neurons for *CG14186* (two-way ANOVA, Šídák’s post-test for for maneb effect, * P <0.05, **** P <0.0001 and for genotype effect in RNAi lines unexposed to maneb, # P <0.05, ## P <0.01, ### P <0.001, n =12-20 brains per genotype) (C) adult-specific panneuronal *Wg* RNAi (via ambient temperature shift 18°C-29°C) results in maneb-induced DA neuron loss (two-way ANOVA, Šídák’s post-test for maneb effect, * P <0.05, and for genotype effect # P <0.05, #### P <0.0001, n =12-15 brains per genotype). Genotypes: control (*elav-GAL4*; *UAS-Dcr2*/*tub-Gal80^ts^*); *Wg* RNAi-1 (*elav-GAL4*; *UAS-Dcr2/ tub-Gal80^ts^*; *Wg-IR* (32994)); *Wg* RNAi-2 (*elav-GAL4*; *tub-Gal80^ts^*; *Wg-IR* (33902)).

### Deregulated Wnt signaling in maneb-related DA neuron vulnerability

Both *fz2* and the mammalian ortholog of *CG14186* (*TMEM237*) have been described to play a role in the highly conserved Wnt signaling pathway. Within the brain, Wnt signaling is critical for neuronal development and synaptic connection formation but is also thought to play a role post-development in the maintenance of synaptic structural changes in response to neuronal activity (21). Toward differentiating developmental vs. non-developmental roles for Wnt signaling in maneb-related neurodegeneration, we attempted to selectively silence this pathway in adult flies via conditional knock-down of the fly Wnt ligand, *wg*. *Gal80^ts^*-based suppression of *wg* RNAi during development was blocked in adult flies after eclosion by an ambient temperature shift from 18°C to 29°C (Methods). Following our standard maneb exposure for 7 days followed by 14 days on regular food under these *GAL4* permissive conditions, we observe significant maneb-induced DA neuron loss in two independent *wg* knock-down lines but not in a control line (Fig. 4C). We additionally observe a slight but significant reduction in DA neurons in one *wg* RNAi line in the absence of maneb. Hence, taken together our data suggest that deregulated Wnt signaling, mediated through knock-down of *fz2* or *wg* and putatively through *CG14186* can promote DA neurodegeneration in flies exposed to maneb.

## DISCUSSION

Genetically tractable model organisms such as *Drosophila* can greatly facilitate efforts to identify genes associated with PD-relevant phenotypes in an unbiased manner (22). Flies have emerged as a powerful model organism for investigating PD and have provided many critical molecular insights into the neurodegenerative process including in response to neurotoxins such as pesticides (23–25). Despite this, few prior studies have investigated neurodegenerative phenotypes in maneb treated flies and those existing studies report mixed results of either no locomotor defects and DA neuron loss in flies exposed to maneb alone for 4 weeks (18), or significant locomotor deficits with subtle whole-body cell death in flies exposed for 15 days (17). Similar modest effects of maneb exposure have typically been reported in mouse models (9, 10). These findings contrast to studies in which flies or mice are exposed to a combination of maneb and another pesticide such as paraquat where more pronounced phenotypes are observed consistent with synergistic effects (9, 10, 18). Notably, these prior fly studies were both conducted in male flies belonging to Canton S (18) or the Harwich strain (17). If gene-environment interactions influence pathologic outcomes, then neurodegenerative phenotypes may be masked in studies of individual isogenic strains. Here, we developed a model of delayed-onset DA neuron loss following maneb exposure and identified quantitatively divergent DA neuron phenotypes across many DGRP fly strains exposed to maneb under tightly controlled conditions. The range of phenotypes observed across different strains argues that genetic architecture does influence maneb sensitivity and supports a role for gene-environment interactions in neurodegeneration. We harnessed this phenotypic divergence in DA neuron viability to perform a genome-wide analysis for candidate associating polymorphisms and through subsequent RNAi studies, identified *fz2* and *CG14186* as modifiers of maneb-related DA neurodegeneration (Fig. 4). *fz2* is a key Wnt ligand receptor orthologous to several human frizzled genes including *FZD5* while *CG14186* is predicted to play a role in cilium assembly and is orthologous to human *TMEM237*. Informatively, both genes and/or their orthologs are implicated in Wnt signaling which is highly conserved between flies and mammals, and we further demonstrate that broad interference of the Wnt pathway via knock-down of the Wnt ligand *wg* promotes DA neuron loss following maneb exposure (Fig. 4C). While the Wnt signaling pathway has a well-established neurodevelopmental role, there is now additional evidence to support its functional involvement in synaptic structure maintenance post-development (21). Further, emerging evidence places deregulated Wnt signaling in the pathogenesis of common neurodegenerative diseases such as Alzheimer’s disease and PD (21, 26). Indeed, reduced Wnt pathway activity was inferred from transcriptional analysis of dopamine neurons isolated from flies exposed to the PD-linked pesticide rotenone (27) and bolstering the pathway via armadillo/β-catenin overexpression was sufficient to prevent rotenone-induced locomotor impairments (27). Hence, deficient Wnt signaling may render DA neurons vulnerable to multiple chemical neurotoxins, although this is an open question that awaits further investigation.

One caveat of our study is the apparent low number of causal genes identified by gene-level RNAi studies, where modifier effects were observed for only 2 out of 16 candidate genes tested (Fig. 4A). Quantitative traits in *Drosophila* are frequently highly polygenic (28) and if this also applies to our maneb DA neuron phenotype, individual candidate gene silencing may simply not be sufficient to manifest frank neuron loss. We did not examine more subtle dopaminergic phenotypes following individual gene knock-down that might potentially illuminate roles for additional candidate genes. There are limitations of our maneb exposure model, however, that may also help explain this result. Chiefly, we do not detect overt DA neuron loss in control strains exposed to 1 mM maneb in our testing protocol (Fig. 4A) which impairs our ability to observe the full range of phenotype modifier effects and especially precludes our ability to observe suppressor effects upon gene knock-down. While this prompted us to narrow our focus to identifying maneb resistance genes that when knocked down precipitate a loss of DA neurons, it limits our effective scope for understanding whether/how genes nominated by GWAS impact maneb-induced neurodegeneration. Conceptually, one way to circumvent this limitation would be to increase the concentration of maneb added to fly food with the goal of promoting DA neuron loss in control genotypes, but since we found that higher maneb concentrations have the undesirable effect of interfering with fly feeding behavior, we opted not to take this approach. Adjusting other parameters of the model, e.g. maneb exposure time or fly age upon initial exposure could also conceivably potentiate phenotypes in control genotypes and in turn enable more comprehensive gene-level testing. These were not tested here but would be important in future studies aimed at refining our maneb exposure model. Despite this limitation, we obtained evidence supporting the contributions of *fz2* and *CG14186* to maneb resistance that led us to elucidate the potential for deregulated Wnt signaling as a possible cause of maneb-related dopaminergic neurodegeneration, thus yielding insight that may inform future PD studies.

Our study extends use of the DGRP to identify genes associated with complex neurodegenerative disease-related traits which has previously included retinal degeneration in retinitis pigmentosa (29), Amyloid β and tau toxicity in a model of Alzheimer’s disease (30) and mutant LRRK2 G2019S-related locomotor deficits in a model of PD (31). Prior studies have also examined the genetic basis of mortality in DGRP strains exposed to the PD-associated pesticide paraquat (32) or to dichlorodiphenyltrichloroethane (DDT) (33), but not in models of PD-related neurodegeneration. Compelling evidence supports a contribution of genetic variants to the risk of PD development (34) and there is also now substantial evidence corroborating a role for gene-environment interactions in neurodegeneration associated with pesticide exposure (35–37). The facile genetics of *Drosophila* coupled to its robust genetic conservation with mammals, short lifespan and ease of implementing precise pesticide exposure presents a unique opportunity to identify genetic factors contributing to pesticide-induced neurodegeneration which can be leveraged to yield novel insight into disease development.

## DATA AVAILABILITY STATEMENT

All DGRP2 lines used in this study are available from the Bloomington *Drosophila* Stock Center at the University of Indiana, Bloomington, IN. File Data S1 contains raw phenotype association data. All DGRP genotype data and SNP location data (Freeze 2.0 calls) are open and can be downloaded from the DGRP data portal (http://dgrp2.gnets.ncsu.edu/data/website/dgrp2.tgeno).

## ACKNOWLEDGMENTS

We thank the Oregon Health and Science University Advanced Light Microscopy Core (RRID:SCR_009961) and Judit Pallos for expert technical assistance. This work was supported by Department of Defense Grant PD200015 to I.M. and made possible in part by the support of Ron and Mary Beamer, whose generosity is deeply appreciated.

## AUTHOR CONTRIBUTIONS

S.V-C. – Conceptualization, Investigation, Formal analysis, Writing manuscript A.A-B. – Investigation, Formal analysis, Writing manuscript I.M. – Conceptualization, Investigation, Formal analysis, Writing manuscript, Supervision, Funding acquisition.

## MATERIALS AND METHODS

### Drosophila stocks and culture

186 DGRP lines used in the study (Table S1) were obtained from the Bloomington *Drosophila* Stock Center as were (stock numbers in parentheses): *TH-GAL4* (8848), *elav-GAL4* (8765); *tub-Gal80^ts^* (7108), RNAi lines: *cv-c* (6403), *Eip74EF* (29353), *Sema-2a* (35432), *Sfp77F* (67376), *Ptp61F* (32426), *cdi* (42568), *kek2* (31874), *CG14186* (29420), *Cad99C* (35037), *fz* (34321), *CadN2* (40889), *Cad87A* (28716), *fz2* (27568), *antp* (64926), *wg* (*wg* RNA-1 (32994), *wg* RNA-2 (33902)), the VALIUM 10/20 vector control *UAS-LUC.VALIUM10* (35788) and *elav-GAL4; UAS-Dcr2* (25750). Additional RNAi lines were obtained from the Vienna *Drosophila* Resource Center: *CG7342* (48001) and *CG10283* (101168). Flies were reared at 25°C/60% relative humidity under a 12 h light-dark cycle on standard cornmeal medium. For *Gal80^ts^* experiments, flies were reared at 18°C/60% relative humidity and aged at 29°C.

### Maneb feeding

Maneb or DMSO control was vigorously stirred into molten standard fly food to a 1 mM final concentration unless otherwise indicated and the food was immediately cooled and stored at 4°C until use and for less than 1 week in total. 50 female adult flies (average age of 2 days-old) from each DGRP line or RNAi cross were collected and transferred to vials containing maneb-supplemented food or vials containing standard food without maneb as control. Flies were housed on maneb-supplemented food for a total of 7 days at 25°C, with a single transfer to fresh maneb food at day 4. Flies were then switched to standard fly food for 14 days with transfer to fresh food every 3-4 days.

### Food intake

Blue dye (erioglaucine disodium salt) was added at 1.5% (w/v) to molten media along with maneb (or DMSO control) at the indicated concentrations just prior to cooling, using previously established procedures (38). Flies (n = 8-10 per condition) were transferred to blue dye-labeled food for 2h then transferred into microtubes for flash freezing in liquid nitrogen. Flies were washed several time in ice-cold water and lysed in ice-cold water. Homogenates were centrifuged at 13,000 rpm for 5 min at 4 °C and resulting supernatants were transferred. Supernatant blue dye concentrations were measured spectrophotometrically at 620 nm (Varioskan Lux). Sample readings were used to calculate average food volume consumed per fly.

### Dopamine neuron immunostaining

Dopamine neuron immunostaining for GWAS and subsequent RNAi experiments were performed as follows: Fly brains (*n* =12-20 per genotype) were collected from female heads that had been pre-fixed in 4% PFA in 0.3% PBS-T for 15 min. Brains were subsequently fixed and permeabilized with 4% PFA in 0.3% PBS-T, pH 7.4 for 20 min at RT, then blocked in 2.5% normal donkey serum for 1 h at room temperature. Brains were incubated in primary anti-TH antibody (1:1000, Immunostar) for three nights on a nutator at 4°C, washed thoroughly and then incubated with Alexa Fluor 488 secondary antibody (1:2000) for 3 nights at 4°C. Brains were again washed extensively before mounting in SlowFade Gold Antifade for imaging. Confocal z-stack images through the entire fly brain were acquired on a Zeiss LSM 900 with a 1 μm slice interval. Dopamine neurons belonging to the protocerebral posterior lateral 1 (PPL1), protocerebral posterior medial 1/2 (PPM1/2), and protocerebral posterior medial 3 (PPM3) clusters were counted and the total from these three clusters was used for genome-wide association analysis.

### Genome-wide association

DA neuron counts for 186 DGRP lines (*n* =12-20 female brains/genotype) exposed to maneb-supplemented food as described above were obtained over a total of 15 experimental batches, with each batch consisting of 8-14 DGRP lines and (except batches 2-5) a previously identified maneb-sensitive line (RAL-911) and a maneb-resistant line (RAL-821) as internal controls that were subsequently used to normalize the data across batches. Residuals for each batch were calculated as the difference between the combined mean of RAL-821 and RAL-911 DA neuron counts in that batch relative to their combined mean overall all batches, and used to mean center raw DA neuron counts for each batch. We estimated broad-sense heritability for DA neuron viability within each of the individual 15 batches in an ANOVA using *α_L_*^2^ / (*α_L_*^2^ + *α_E_*^2^), where *α_L_*^2^ represents the between-line variance of raw DA neuron counts and *α_E_*^2^ is the within-line variance. Overall mean heritability was then calculated across all 15 individual batch estimates. Batch-corrected DA neuron count data (or raw DA neuron count data for batches 2-5) consisting of line means for all 186 DGRP lines (Table S1) were uploaded to the DGRP portal (http://dgrp.gnets.ncsu.edu/) which uses PLINK, FaST-LMM, SnpEff and R to perform genome-wide association analysis using the following linear mixed model which controls for population structure and relatedness among lines: *y* = X*b* + Z*u* + *e*, where *y* is the vector of DA neuron counts adjusted for *Wolbachia* and five major inversions, X is the design matrix of fixed SNP effect *b*, Z is the design matrix for the random polygenic effect *u*, and *e* is the residual error. The vector of randem effects *u* has a covariance matrix A*σ*^2^, in which *σ*^2^ is the polygenic variance component. SNPs and other variants are filtered for minor allele frequency (≥ 0.05), missingness (<30%) and non-biallelic sites are removed. A total of 1,889,672 variants were included in the genome-wide analysis which regressed DA neuron counts for 186 DGRP lines onto each variant, the vast majority of which are SNP (Data S1). Q-Q plot analysis supported normally distributed data (Fig. 3C) and covariate analysis indicated that *Wolbachia* infection status and five major chromosomal inversions harbored by the DGRP are not significant phenotype covariates (Table S3). Pairwise linkage disequilibrium (LD) analysis on the 19 highest ranking SNPs (linear mixed model-derived *P* <10^−6^) demonstrates that SNPs in LD (*r^2^* ≥0.5) are all within the same gene boundaries, thus consistent with overall low LD (Figure 3B). The BDGP R54/dm3 genome build was used to identify candidate associating genes from SNP coordinates (freeze 2.0 calls), whereby a SNP was assigned to a gene if it was within that gene’s transcription boundaries ± 1 kb. Genes nominated from SNPs associating at the linear mixed model-derived *P* <10^−5^ threshold (Table S2) were analyzed using the PANTHER 19.0 database using GO biological process-complete enrichment against all *Drosophila* genes in the database and a Fisher’s exact test (Table 2).

### Gene-level RNAi testing

For RNAi screening, males from 16 RNAi transgenic lines (10 top association genes and 6 cadherin/Wnt signaling genes identified by GO analysis) or the VALIUM10/20 vector control line were mated to female flies harboring *TH-GAL4* in a single experimental batch. *TH-GAL4>UAS-RNAi* female progeny (50 per genotype) were collected and exposed to maneb-supplemented food as described above. At the 21-day timepoint, flies (n =12-20) were processed for dopamine neuron immunostaining as described above. For the adult-specific *wg* RNAi experiment, we generated *tub-GAL80^ts^*; *wg RNAi* (32994 or 33902) double transgenic lines and crossed males from these lines to *elavC155-GAL4*; *UAS-Dcr2* or to *elav-GAL4* females. A control genotype was obtained from crossing *tub-GAL80^ts^* males to *elavC155-GAL4*; *UAS-Dcr2* females. Flies were maintained at 18°C throughout progeny development to suppress GAL4, then female progeny (50 per genotype) were collected and switched to 29°C prior to maneb exposure and maintained at 29°C throughout the 21 day maneb on/off exposure protocol described above.

### Statistical analysis

Data were analyzed by Student’s t-test, ANOVA or two-way ANOVA with correction for multiple comparisons unless otherwise noted and *P* <0.05 was considered statistically significant. Data are presented as mean ± SEM and individual data points are shown in graphs for datasets with n σ; 12.

## Notes

### Competing Interest Statement

The authors have declared no competing interest.

