## Supplemental Tables S1-S3 for "Genetic basis of maneb-induced dopaminergic neurodegeneration in *Drosophila*"

**Table S1.** DGRP lines, mean dopamine neuron counts, strain *Wolbachia* infection status and survivorship at the end of 21-day maneb exposure plus aging period.

| **DGRP line number** | **DA neurons (mean)** | ***Wolbachia* infection status** | **Survival at 21 days (%)** |
| --- | --- | --- | --- |
| 21 | 50.3 | Y | 100 |
| 26 | 45.5 | N | 100 |
| 28 | 47.6 | N | 100 |
| 31 | 47.7 | N | 96 |
| 32 | 46.6 | N | 100 |
| 40 | 49.1 | Y | 96 |
| 41 | 47.3 | N | 96 |
| 42 | 48.8 | N | 100 |
| 45 | 44.6 | N | 98 |
| 48 | 50.4 | Y | 100 |
| 57 | 44.5 | N | 98 |
| 59 | 47.6 | N | 100 |
| 69 | 47.9 | Y | 92 |
| 73 | 50.8 | Y | 82 |
| 75 | 48.4 | Y | 98 |
| 83 | 41.2 | N | 94 |
| 85 | 54.1 | N | 100 |
| 88 | 49.3 | N | 100 |
| 91 | 46.0 | N | 91 |
| 93 | 48.7 | N | 100 |
| 100 | 49.2 | Y | 98 |
| 101 | 48.4 | N | 100 |
| 105 | 46.8 | N | 100 |
| 109 | 47.0 | N | 100 |
| 129 | 45.9 | N | 98 |
| 136 | 52.5 | Y | 98 |
| 142 | 50.6 | Y | 98 |
| 149 | 42.6 | Y | 100 |
| 153 | 43.2 | Y | 86 |
| 161 | 46.8 | N | 94 |
| 177 | 40.8 | N | 65 |
| 181 | 48.7 | Y | 98 |
| 189 | 49.7 | Y | 98 |
| **DGRP line number** | **DA neurons (mean)** | ***Wolbachia* infection status** | **Survival at 21 days (%)** |
| 195 | 50.2 | N | 96 |
| 208 | 47.7 | N | 100 |
| 217 | 47.3 | N | 96 |
| 227 | 47.0 | Y | 94 |
| 228 | 48.2 | N | 100 |
| 229 | 47.9 | N | 94 |
| 235 | 50.2 | N | 100 |
| 239 | 46.3 | N | 54 |
| 256 | 48.0 | Y | 92 |
| 280 | 46.4 | Y | 100 |
| 287 | 48.3 | Y | 100 |
| 301 | 45.4 | N | 96 |
| 303 | 45.1 | N | 100 |
| 304 | 51.7 | Y | 100 |
| 306 | 43.9 | Y | 100 |
| 307 | 48.2 | N | 84 |
| 309 | 46.2 | N | 98 |
| 310 | 48.5 | Y | 96 |
| 313 | 53.2 | N | 100 |
| 315 | 43.5 | N | 92 |
| 317 | 44.1 | Y | 100 |
| 318 | 47.9 | Y | 100 |
| 319 | 45.5 | Y | 100 |
| 320 | 46.6 | Y | 100 |
| 321 | 46.8 | Y | 100 |
| 324 | 48.5 | N | 100 |
| 332 | 44.2 | N | 100 |
| 335 | 51.7 | Y | 90 |
| 336 | 46.4 | Y | 98 |
| 338 | 48.4 | Y | 98 |
| 340 | 46.7 | Y | 98 |
| 348 | 47.1 | N | 94 |
| 350 | 49.3 | N | 100 |
| 352 | 43.2 | Y | 94 |
| 354 | 47.7 | N | 100 |
| 355 | 45.7 | Y | 98 |
| 356 | 46.8 | Y | 65 |
| **DGRP line number** | **DA neurons (mean)** | ***Wolbachia* infection status** | **Survival at 21 days (%)** |
| 357 | 46.4 | N | 96 |
| 358 | 46.4 | N | 100 |
| 359 | 50.4 | N | 100 |
| 360 | 50.3 | Y | 99 |
| 361 | 51.8 | Y | 100 |
| 362 | 46.7 | Y | 100 |
| 365 | 47.1 | Y | 100 |
| 370 | 47.5 | Y | 94 |
| 371 | 46.2 | N | 98 |
| 373 | 53.2 | N | 100 |
| 374 | 45.3 | Y | 100 |
| 375 | 48.4 | N | 98 |
| 377 | 48.2 | N | 100 |
| 379 | 46.6 | N | 94 |
| 380 | 42.3 | Y | 98 |
| 381 | 46.3 | N | 98 |
| 383 | 48.2 | Y | 92 |
| 385 | 47.7 | N | 100 |
| 386 | 50.4 | N | 98 |
| 390 | 48.7 | N | 52 |
| 391 | 49.6 | N | 98 |
| 392 | 44.0 | N | 100 |
| 395 | 47.1 | N | 90 |
| 397 | 47.2 | Y | 98 |
| 399 | 48.1 | N | 100 |
| 405 | 46.8 | Y | 100 |
| 406 | 46.6 | N | 98 |
| 426 | 43.5 | N | 88 |
| 427 | 48.5 | N | 100 |
| 437 | 45.4 | N | 98 |
| 439 | 52.1 | N | 100 |
| 440 | 46.0 | Y | 98 |
| 441 | 46.4 | Y | 100 |
| 443 | 49.4 | N | 92 |
| 461 | 44.9 | Y | 98 |
| 486 | 49.5 | Y | 94 |
| 491 | 46.4 | N | 100 |
| **DGRP line number** | **DA neurons (mean)** | ***Wolbachia* infection status** | **Survival at 21 days (%)** |
| 492 | 47.0 | N | 100 |
| 502 | 52.3 | N | 94 |
| 505 | 43.7 | Y | 100 |
| 508 | 46.9 | N | 96 |
| 509 | 45.6 | N | 96 |
| 513 | 50.6 | Y | 100 |
| 517 | 49.4 | N | 100 |
| 528 | 49.1 | Y | 96 |
| 530 | 51.3 | Y | 100 |
| 531 | 47.9 | Y | 92 |
| 535 | 47.1 | Y | 96 |
| 551 | 51.2 | Y | 94 |
| 555 | 45.5 | Y | 100 |
| 559 | 46.4 | N | 100 |
| 563 | 47.6 | N | 98 |
| 566 | 48.4 | N | 100 |
| 584 | 45.4 | Y | 100 |
| 589 | 52.2 | Y | 96 |
| 595 | 44.5 | Y | 90 |
| 596 | 43.9 | N | 55 |
| 627 | 50.5 | N | 100 |
| 630 | 50.4 | N | 98 |
| 634 | 42.9 | Y | 100 |
| 639 | 50.1 | Y | 96 |
| 646 | 49.8 | Y | 76 |
| 703 | 50.2 | N | 98 |
| 705 | 47.8 | Y | 100 |
| 707 | 48.8 | Y | 100 |
| 712 | 49.2 | Y | 100 |
| 714 | 48.7 | N | 70 |
| 716 | 46.9 | Y | 98 |
| 721 | 52.7 | Y | 100 |
| 730 | 47.9 | Y | 96 |
| 732 | 46.5 | N | 100 |
| 737 | 46.3 | Y | 98 |
| 738 | 50.8 | Y | 100 |
| 748 | 40.8 | Y | 100 |
| **DGRP line number** | **DA neurons (mean)** | ***Wolbachia* infection status** | **Survival at 21 days (%)** |
| 757 | 47.4 | N | 67 |
| 761 | 50.8 | Y | 100 |
| 765 | 50.1 | N | 82 |
| 774 | 46.5 | N | 100 |
| 776 | 49.4 | Y | 100 |
| 783 | 46.6 | Y | 96 |
| 786 | 46.9 | Y | 98 |
| 787 | 46.3 | Y | 100 |
| 790 | 44.7 | Y | 98 |
| 796 | 48.5 | Y | 98 |
| 799 | 45.3 | N | 91 |
| 801 | 44.6 | Y | 100 |
| 802 | 47.6 | Y | 92 |
| 804 | 49.8 | Y | 88 |
| 808 | 49.0 | N | 100 |
| 810 | 48.6 | N | 100 |
| 812 | 42.6 | N | 100 |
| 818 | 48.8 | Y | 98 |
| 819 | 43.4 | Y | 78 |
| 821 | 48.9 | Y | 100 |
| 822 | 50.1 | Y | 98 |
| 832 | 44.0 | Y | 98 |
| 837 | 46.6 | Y | 100 |
| 843 | 42.7 | N | 94 |
| 850 | 48.5 | Y | 96 |
| 852 | 48.7 | Y | 100 |
| 853 | 49.2 | Y | 98 |
| 855 | 51.3 | Y | 100 |
| 857 | 49.2 | N | 92 |
| 859 | 47.7 | Y | 100 |
| 861 | 49.7 | Y | 94 |
| 879 | 47.4 | Y | 100 |
| 882 | 48.7 | Y | 100 |
| 884 | 47.5 | Y | 84 |
| 890 | 48.5 | Y | 96 |
| 892 | 47.6 | Y | 100 |
| 897 | 43.5 | Y | 100 |
| **DGRP line number** | **DA neurons (mean)** | ***Wolbachia* infection status** | **Survival at 21 days (%)** |
| 900 | 45.1 | N | 94 |
| 907 | 46.6 | N | 98 |
| 908 | 42.9 | N | 90 |
| 911 | 42.0 | N | 65 |
| 913 | 46.9 | Y | 100 |

**Table S2.** Extended list of associating SNP (P<10^-5^). MAF, minor allele frequence. Gene is the gene boundaries ± 1 kb where each SNP is located.

| **SNP ID** | **MAF** | **Minor**  **Allele**  **Count** | **Major**  **Allele**  **Count** | **Mixed**  **model**  ***P*-value** | **Gene** |
| --- | --- | --- | --- | --- | --- |
| 3R_10296407_SNP | 0.1337 | 23 | 149 | 4.26E-07 | *cv-c* |
| 2L_18476602_SNP | 0.1011 | 18 | 160 | 6.49E-07 | *CG10283* |
| 3L_17578294_SNP | 0.0565 | 10 | 167 | 9.08E-07 | *Eip74EF* |
| 3R_10294090_SNP | 0.05525 | 10 | 171 | 1.07E-06 | *cv-c* |
| 3R_10299555_SNP | 0.05028 | 9 | 170 | 1.13E-06 | *cv-c* |
| 2L_18476619_SNP | 0.1061 | 19 | 160 | 1.73E-06 | *CG10283* |
| 3L_20894399_SNP | 0.1792 | 31 | 142 | 3.19E-06 | *Sfp77F* |
| 3R_10296305_SNP | 0.1364 | 24 | 152 | 4.01E-06 | *cv-c* |
| 2R_12396038_SNP | 0.2994 | 50 | 117 | 4.03E-06 | *Sema-2a* |
| 3L_1356451_SNP | 0.1196 | 22 | 162 | 4.39E-06 | *Ptp61F* |
| 3R_16888532_SNP | 0.1099 | 20 | 162 | 4.84E-06 | *Cortactin* |
| 3R_14904153_SNP | 0.06593 | 12 | 170 | 6.41E-06 | *cdi* |
| 3R_14904152_SNP | 0.06593 | 12 | 170 | 7.38E-06 | *cdi* |
| 2L_11613526_SNP | 0.09827 | 17 | 156 | 7.56E-06 | *kek2* |
| 3L_20094140_SNP | 0.1871 | 32 | 139 | 9.24E-06 | *CG14186* |
| 3R_15432051_SNP | 0.4451 | 73 | 91 | 9.40E-06 | *CG7342* |
| 3R_10298709_SNP | 0.1552 | 27 | 147 | 9.54E-06 | *cv-c* |
| 2R_12396133_SNP | 0.3876 | 69 | 109 | 1.06E-05 | *Sema-2a* |
| 3R_10298462_SNP | 0.1471 | 25 | 145 | 1.10E-05 | *cv-c* |
| 3L_13321889_SNP | 0.08065 | 15 | 171 | 1.18E-05 | *CG14113* |
| 2L_19857167_SNP | 0.2907 | 50 | 122 | 1.23E-05 | *sick* |
| 3R_10299135_SNP | 0.05587 | 10 | 169 | 1.38E-05 | *cv-c* |
| 3L_20897935_SNP | 0.4371 | 73 | 94 | 1.78E-05 | *CG11458* |
| 3L_14904022_SNP | 0.1742 | 27 | 128 | 1.80E-05 | *DCX-EMAP* |
| 2L_18813454_SNP | 0.06044 | 11 | 171 | 1.91E-05 | *CG42305* |
| 3R_8873317_SNP | 0.3529 | 60 | 110 | 1.98E-05 | *CG31157* |
| 2R_19287782_SNP | 0.2707 | 49 | 132 | 2.02E-05 | *CG3520* |
| 3R_12615678_SNP | 0.3314 | 58 | 117 | 2.02E-05 | *Glut-3* |
| 3R_15432152_SNP | 0.4405 | 74 | 94 | 2.09E-05 | *CG7342* |
| 2L_6143751_SNP | 0.2976 | 50 | 118 | 2.16E-05 | *CG13991* |
| 2R_7986403_SNP | 0.3022 | 55 | 127 | 2.30E-05 | *jeb* |
| 3L_13321857_SNP | 0.05495 | 10 | 172 | 2.36E-05 | *CG14113* |
| **SNP ID** | **MAF** | **Minor**  **Allele**  **Count** | **Major**  **Allele**  **Count** | **Mixed**  **model**  **P-value** | **Gene** |
| 3R_17994440_SNP | 0.0791 | 14 | 163 | 2.40E-05 | *SKIP* |
| 3L_3004825_SNP | 0.06486 | 12 | 173 | 2.42E-05 | *FMRaR* |
| 2L_19857160_SNP | 0.2965 | 51 | 121 | 2.45E-05 | *sick* |
| 3L_19192232_SNP | 0.1356 | 24 | 153 | 2.48E-05 | *fz2* |
| X_17889690_SNP | 0.4111 | 74 | 106 | 2.60E-05 | *Sh* |
| X_10613019_SNP | 0.453 | 82 | 99 | 2.62E-05 | *stx* |
| 3R_25483389_SNP | 0.1017 | 18 | 159 | 2.86E-05 | *Dop1R2* |
| 3R_5447409_SNP | 0.08824 | 15 | 155 | 3.17E-05 | *CG8312* |
| 2L_10902703_SNP | 0.1461 | 26 | 152 | 3.25E-05 | *CG14070* |
| 3L_2908300_SNP | 0.1236 | 22 | 156 | 3.37E-05 | *shab* |
| 3L_19190825_SNP | 0.1742 | 31 | 147 | 3.49E-05 | *fz2* |
| 3L_14330568_SNP | 0.3372 | 58 | 114 | 3.64E-05 | *fz* |
| 3L_3133505_SNP | 0.3039 | 55 | 126 | 3.67E-05 | *CG11537* |
| 3L_11093584_SNP | 0.2044 | 37 | 144 | 3.71E-05 | *APP-BP1* |
| X_10613158_SNP | 0.4556 | 82 | 98 | 3.71E-05 | *stx* |
| 2R_7403101_SNP | 0.2453 | 39 | 120 | 3.82E-05 | *CG30034* |
| 3R_11209466_SNP | 0.2278 | 41 | 139 | 3.84E-05 | *Rbp* |
| 3R_10298469_SNP | 0.1529 | 26 | 144 | 3.85E-05 | *cv-c* |
| 3L_8762342_SNP | 0.05914 | 11 | 175 | 3.95E-05 | *Fhos* |
| 2L_21358911_SNP | 0.08 | 14 | 161 | 4.03E-05 | *Tsp39D* |
| 3L_11183025_SNP | 0.1326 | 24 | 157 | 4.03E-05 | *GlcAT-P* |
| 3L_3133827_SNP | 0.2278 | 41 | 139 | 4.05E-05 | *CG11537* |
| X_10613157_SNP | 0.4505 | 82 | 100 | 4.05E-05 | *stx* |
| 3R_17169476_SNP | 0.06111 | 11 | 169 | 4.06E-05 | *aus* |
| X_17146922_SNP | 0.05325 | 9 | 160 | 4.23E-05 | *f* |
| 3R_17430744_SNP | 0.05682 | 10 | 166 | 4.30E-05 | *InR* |
| 3L_2908286_SNP | 0.2047 | 35 | 136 | 4.35E-05 | *Shab* |
| 2L_18819809_SNP | 0.05 | 9 | 171 | 4.43E-05 | *Ugt36F1* |
| 3L_20097398_SNP | 0.2126 | 37 | 137 | 4.48E-05 | *CG14186* |
| 3L_2842577_SNP | 0.05978 | 11 | 173 | 4.50E-05 | *Tet* |
| 3R_8911066_SNP | 0.1011 | 18 | 160 | 4.64E-05 | *MIP20204* |
| X_17146949_SNP | 0.05172 | 9 | 165 | 4.76E-05 | *f* |
| X_10613015_MNP | 0.4481 | 82 | 101 | 4.79E-05 | *stx* |
| 3L_8759522_SNP | 0.0618 | 11 | 167 | 4.87E-05 | *Fhos* |
| X_5340317_SNP | 0.3239 | 57 | 119 | 4.97E-05 | *SPR* |
| 3L_14330327_SNP | 0.3462 | 63 | 119 | 5.05E-05 | *fz* |
| 2L_19857135_SNP | 0.4639 | 77 | 89 | 5.10E-05 | *sick* |
| **SNP ID** | **MAF** | **Minor**  **Allele**  **Count** | **Major**  **Allele**  **Count** | **Mixed**  **model**  **P-value** | **Gene** |
| 3R_5563006_SNP | 0.07865 | 14 | 164 | 5.11E-05 | *side-VII* |
| 2R_11735502_SNP | 0.264 | 47 | 131 | 5.29E-05 | *tun* |
| 3L_579756_SNP | 0.3812 | 69 | 112 | 5.30E-05 | *Hipk* |
| X_12939649_SNP | 0.3646 | 66 | 115 | 5.31E-05 | *rad* |
| X_6933740_SNP | 0.4973 | 92 | 93 | 5.31E-05 | *CG12541* |
| 2R_11694842_SNP | 0.435 | 77 | 100 | 5.38E-05 | *Zasp52* |
| 2R_18710365_SNP | 0.25 | 45 | 135 | 5.51E-05 | *CG42284* |
| X_17391235_SNP | 0.324 | 58 | 121 | 5.56E-05 | *CG43658* |
| 3L_2757984_SNP | 0.1768 | 32 | 149 | 5.64E-05 | *Mrtf* |
| X_10613125_SNP | 0.4586 | 83 | 98 | 5.73E-05 | *stx* |
| 3R_17498869_SNP | 0.08046 | 14 | 160 | 5.78E-05 | *CG31176* |
| X_10613097_SNP | 0.453 | 82 | 99 | 6.01E-05 | *stx* |
| 3L_15943976_SNP | 0.1923 | 35 | 147 | 6.10E-05 | *Pka-C3* |
| 2L_19864680_SNP | 0.2189 | 37 | 132 | 6.28E-05 | *sick* |
| 2L_16412303_SNP | 0.3964 | 67 | 102 | 6.31E-05 | *BG:DS02780.1* |
| X_10613013_SNP | 0.4426 | 81 | 102 | 6.44E-05 | *stx* |
| 3L_3133555_SNP | 0.221 | 40 | 141 | 6.45E-05 | *CG11537* |
| 3L_14904021_SNP | 0.122 | 20 | 144 | 6.59E-05 | *DCX-EMAP* |
| X_5605787_SNP | 0.453 | 82 | 99 | 6.61E-05 | *CG33080* |
| 2R_19896049_SNP | 0.05405 | 10 | 175 | 6.77E-05 | *Unc-89* |
| 3L_3134029_SNP | 0.1593 | 29 | 153 | 6.77E-05 | *CG11537* |
| 2L_16412312_SNP | 0.407 | 70 | 102 | 6.79E-05 | *BG:DS02780.1* |
| 3L_3014623_SNP | 0.173 | 32 | 153 | 6.89E-05 | *CG32488* |
| 2R_7992945_SNP | 0.3466 | 61 | 115 | 6.91E-05 | *jeb* |
| X_10613133_SNP | 0.4511 | 83 | 101 | 6.93E-05 | *stx* |
| 3L_20889901_SNP | 0.2882 | 49 | 121 | 6.95E-05 | *Sfp77F* |
| 2L_3455643_SNP | 0.2023 | 35 | 138 | 7.00E-05 | *Cog3* |
| X_1120169_SNP | 0.1593 | 29 | 153 | 7.03E-05 | *Motor* |
| X_6179699_SNP | 0.1421 | 26 | 157 | 7.08E-05 | *Top3beta* |
| X_10613026_SNP | 0.4432 | 82 | 103 | 7.08E-05 | *stx* |
| 2L_3454743_SNP | 0.2609 | 42 | 119 | 7.16E-05 | *Cog3* |
| 2R_16422217_SNP | 0.1517 | 27 | 151 | 7.19E-05 | *Bbd* |
| 2R_11796133_SNP | 0.2346 | 42 | 137 | 7.30E-05 | *sli* |
| 3L_19195311_SNP | 0.06145 | 11 | 168 | 7.35E-05 | *fz2* |
| 2L_18476867_SNP | 0.07182 | 13 | 168 | 7.40E-05 | *CG10283* |
| 3R_2802793_SNP | 0.2197 | 38 | 135 | 7.48E-05 | *Antp* |
| 2L_4008120_SNP | 0.1824 | 31 | 139 | 7.49E-05 | *CG10039* |
| **SNP ID** | **MAF** | **Minor**  **Allele**  **Count** | **Major**  **Allele**  **Count** | **Mixed**  **model**  **P-value** | **Gene** |
| 2R_13018698_SNP | 0.06522 | 12 | 172 | 7.55E-05 | *CG45273* |
| 3L_2755846_SNP | 0.2391 | 44 | 140 | 7.57E-05 | *Mrtf* |
| 2R_3355650_SNP | 0.3918 | 67 | 104 | 7.59E-05 | *Aldh-III* |
| 2R_19502307_SNP | 0.06145 | 11 | 168 | 7.61E-05 | *Eglp2* |
| X_10613048_SNP | 0.4511 | 83 | 101 | 7.71E-05 | *stx* |
| 3R_19448792_SNP | 0.05464 | 10 | 173 | 7.74E-05 | *CG31145* |
| 2R_13018775_SNP | 0.06452 | 12 | 174 | 7.80E-05 | *CG45273* |
| 2L_19631006_SNP | 0.1582 | 28 | 149 | 7.86E-05 | *Lar* |
| 2R_14915139_SNP | 0.309 | 55 | 123 | 7.86E-05 | *5-HT1B* |
| 2L_22124577_SNP | 0.3091 | 51 | 114 | 7.87E-05 | *CG3651* |
| 2R_13018691_SNP | 0.06486 | 12 | 173 | 8.10E-05 | *CG45273* |
| X_15360491_SNP | 0.1639 | 30 | 153 | 8.19E-05 | *CG6299* |
| 3L_18626730_SNP | 0.07263 | 13 | 166 | 8.26E-05 | *PT-75D* |
| 3R_21491664_SNP | 0.1802 | 31 | 141 | 8.32E-05 | *Lgr3* |
| 2L_18915465_SNP | 0.1067 | 19 | 159 | 8.39E-05 | *ssp3* |
| 3L_2750780_SNP | 0.2044 | 37 | 144 | 8.41E-05 | *Mrtf* |
| 2L_17808538_SNP | 0.2486 | 43 | 130 | 8.44E-05 | *CadN2* |
| 3R_7763244_SNP | 0.1322 | 23 | 151 | 8.45E-05 | *Cad87A* |
| 3R_22422124_SNP | 0.06667 | 12 | 168 | 8.48E-05 | *scrib* |
| 3L_6506214_SNP | 0.4765 | 81 | 89 | 8.71E-05 | *sfl* |
| 3R_24065322_SNP | 0.06536 | 10 | 143 | 8.80E-05 | *CG34354* |
| 3R_25672141_SNP | 0.1695 | 30 | 147 | 8.81E-05 | *Cad99C* |
| 3L_21412875_SNP | 0.05917 | 10 | 159 | 8.83E-05 | *rgn* |
| 2R_7953435_SNP | 0.4831 | 86 | 92 | 8.87E-05 | *CG8888* |
| 3L_19194617_SNP | 0.1461 | 26 | 152 | 8.97E-05 | *fz2* |
| 3L_19190762_SNP | 0.1348 | 24 | 154 | 8.97E-05 | *fz2* |
| 2L_1085727_MNP | 0.2727 | 48 | 128 | 9.07E-05 | *CG4629* |
| X_10613053_MNP | 0.4481 | 82 | 101 | 9.10E-05 | *stx* |
| 3L_2842279_SNP | 0.05525 | 10 | 171 | 9.10E-05 | *Tet* |
| 2R_7953497_SNP | 0.3889 | 70 | 110 | 9.16E-05 | *CG8888* |
| 2R_10205440_SNP | 0.0838 | 15 | 164 | 9.38E-05 | *Shrm* |
| 2L_12597209_SNP | 0.1067 | 19 | 159 | 9.41E-05 | *nub* |
| 3L_5005815_SNP | 0.06044 | 11 | 171 | 9.46E-05 | *Con* |
| 2L_2459998_SNP | 0.3779 | 65 | 107 | 9.72E-05 | *dpp* |
| 3R_8873700_SNP | 0.4471 | 76 | 94 | 9.73E-05 | *CG31157* |
| 2L_19870642_SNP | 0.2321 | 39 | 129 | 9.78E-05 | *sick* |
| 2L_3454725_SNP | 0.2959 | 50 | 119 | 9.92E-05 | *Cog3* |
| **SNP ID** | **MAF** | **Minor**  **Allele**  **Count** | **Major**  **Allele**  **Count** | **Mixed**  **model**  **P-value** | **Gene** |
| 3R_9768093_SNP | 0.125 | 22 | 154 | 9.94E-05 | *Art9* |
| 3R_10287583_SNP | 0.05028 | 9 | 170 | 9.95E-05 | *cv-c* |
| 3L_12426677_SNP | 0.07527 | 14 | 172 | 9.98E-05 | *toe* |

**Table S3**. Type III ANOVA table for *Wolbachia* and inversion covariates.

| **Factor** | **Df** | **Sum Sq** | **RSS** | **AIC** | **F value** | **Pr(>F)** |
| --- | --- | --- | --- | --- | --- | --- |
| *Wolbachia* | 1 | 4.756 | 1155.4 | 361.72 | 0.7192 | 0.3976 |
| In_2L_t | 2 | 5.949 | 1156.6 | 359.91 | 0.4498 | 0.6384 |
| In_2R_NS | 2 | 0.988 | 1151.6 | 359.11 | 0.0747 | 0.9281 |
| In_3R_P | 2 | 14.660 | 1165.3 | 361.3 | 1.1084 | 0.3324 |
| In_3R_K | 2 | 5.690 | 1156.3 | 359.87 | 0.4302 | 0.6511 |
| In_3R_Mo | 2 | 20.542 | 1171.2 | 362.24 | 1.5532 | 0.2145 |
